# Non-Covalently Associated Streptavidin Multi-Arm Nanohubs Exhibit Mechanical and Thermal Stability in Protein-Network Materials

**DOI:** 10.1101/2022.05.09.491252

**Authors:** David S. Knoff, Samuel Kim, Kareen A. Fajardo Cortes, Jocelyne Rivera, Dallas Altamirano, Minkyu Kim

**Affiliations:** Department of Biomedical Engineering, University of Arizona, Tucson, AZ 85721; Department of Materials Science and Engineering, University of Arizona, Tucson, AZ 85721; BIO5 Institute, University of Arizona, Tucson, AZ 85719

**Keywords:** Molecular self-assembly, Protein cross-linkers, Non-covalent interactions, Artificial protein design, Hierarchical assembly, Protein-network, Hydrogel

## Abstract

Constructing protein-network materials that exhibit physicochemical and mechanical properties of individual protein constituents requires molecular cross-linkers with specificity and stability. A well-known example involves specific chemical fusion of a four-arm polyethylene glycol (tetra-PEG) to desired proteins with secondary cross-linkers. However, it is necessary to investigate tetra-PEG like biomolecular cross-linkers that are genetically fused to the proteins, simplifying synthesis by removing additional conjugation and purification steps. Non-covalently, self-associating, streptavidin homotetramer is a viable, biomolecular alternative to tetra-PEG. Here, a multi-arm streptavidin design is characterized as a protein-network material platform using various secondary, biomolecular cross-linkers, such as high-affinity physical (i.e., non-covalent), transient physical, spontaneous chemical (i.e., covalent), or stimuli-induced chemical cross-linkers. Stimuli-induced, chemical cross-linkers fused to multi-arm streptavidin nanohubs provide sufficient diffusion prior to initiating permanent covalent bonds, allowing proper characterization of streptavidin nanohubs. Surprisingly, non-covalently associated streptavidin nanohubs exhibit extreme stability which translates into material properties that resemble hydrogels formed by chemical bonds even at high temperatures. Therefore, this study not only establishes that the streptavidin nanohub is an ideal multi-arm biopolymer precursor but also provides valuable guidance for designing self-assembling nanostructured molecular networks that can properly harness the extraordinary properties of protein-based building blocks.

## Introduction

Functional proteins and molecular cross-linkers constitute protein-network materials, ubiquitous in nature, such as cytoskeleton^1, 2^ and extracellular matrix.^3, 4^ Understanding bulk physicochemical, biological, and single-molecule mechanical properties of functional proteins provides the opportunity to develop synthetic protein-network materials that mimic native biology for tissue-engineered scaffolds,^5^ and further implement desired properties, such as resilience,^6, 7^ passive elasticity of muscle,^8^ strength,^9^ and adhesion in aqueous environments.^10, 11^ Diverse synthetic and biological molecular cross-linkers are available to connect and exhibit physicochemical and biological properties of proteins in protein-network systems.^7, 12–15^ However, to exhibit bulk mechanical properties of protein-network materials, depending on constitutive mechanical proteins, selecting molecular cross-linkers requires more stringent conditions, such as specificity, functionality, and strength.^12^ For instance, individual mechanical proteins exhibit their properties when external stresses stretch and unfold the protein via each terminus,^8, 15–18^ necessitating high cross-linker specificity for crosslinking proteins end-to-end within the network. When cross-linkers are located at both termini of each protein, at least one cross-linker should have the functionality to connect with at least two other cross-linkers from separate functional proteins (*f* ≥ 3) to prepare a three-dimensional protein-network system. Furthermore, cross-linkers need to remain stable and intact to transmit the applied external stress across all protein strands in the protein network. Currently, such well-defined molecular cross-linkers that concurrently have high specificity and stability as well as the functionality within protein-network materials are limited.

A molecular cross-linker that meets these requirements is a four-arm polyethylene glycol (tetra-PEG)^19^ that has been chemically fused with specificity to proteins of interest with secondary cross-linkers at the other ends to form protein-network materials.^12^ Tetra-PEG, which has a crosslinking functionality of four (*f* = 4), represents a trending strategy to develop star-like precursors that form polymer-network materials.^12, 20, 21^ A major disadvantage of this approach to fabricate protein-network materials is the conjugation and purification steps required to chemically fuse tetra-PEG to proteins, which increases the complexity of synthesis and processing, and can reduce yields of the star-like precursors. This leads to the investigation of tetra-PEG-like protein cross-linkers that can be genetically fused to proteins of interest, simplifying multi-arm protein synthesis as well as improving customizability and promoting the specific incorporation of functional proteins into strands of protein-network materials via genetic and protein engineering technology.^22^

Secondary or tertiary structures of proteins that self-recognize and non-covalently self-associate each other, constituting quaternary or oligomeric structures that have been used as biomolecular cross-linkers. By genetically fusing the cross-linkers to proteins of interest using recombinant DNA technology, biosynthesized artificial proteins can construct predictable macromolecular structures,^23–26^ such as multi-arm star-like precursors similar to tetra-PEG cross-linkers (*f* ≥ 3).^23^ Furthermore, this approach enables the fabrication of protein-network materials with cross-linkers containing tailored binding affinity, modulating the intrinsic nature of protein associations and strands with customized length and mechanics.^27–31^ Despite these findings, the lack of cross-linker stability due to non-covalent interactions requires the investigation of strong and specific biomolecular cross-linkers to serve as multi-arm precursors that produce stable protein-network materials.

Well-known for having the strongest non-covalent interactions in nature with its ligand, biotin (*K_d_* = 10^-14^ ∼10^-16^ M),^32, 33^ streptavidin (SAv) has been investigated for diverse biotechnology applications,^34^ including binding assays, and purification strategies. Moreover, bulk biochemical study recognized that SAv forms thermally stable tetramers that can resist heat up to 75 °C and chemically stable against protein denaturants, such as 6M GuHCl.^35^ It is also identified that SAv tetramers are mechanically stable, specifically at the monomer-monomer interface where AFM-based single-molecular force-spectroscopy study showed that this interface is four times stronger than the dimer-dimer interface.^23^ Furthermore, these information has been the foundation to develop stable and precise nano-assemblies by applying the concept that SAv monomers naturally self-assemble into homotetramers with high affinity.^36^ This indicates that SAv can be a viable strong and specific biomolecular cross-linker. When genetically fusing proteins of interest to one end of SAv monomers, biosynthesized artificial proteins will self-recognize and self-associate into a stable four-arm star-like precursor (*f* = 4), tetra-SAv (**Figure 1**).^23^

**Figure 1.**
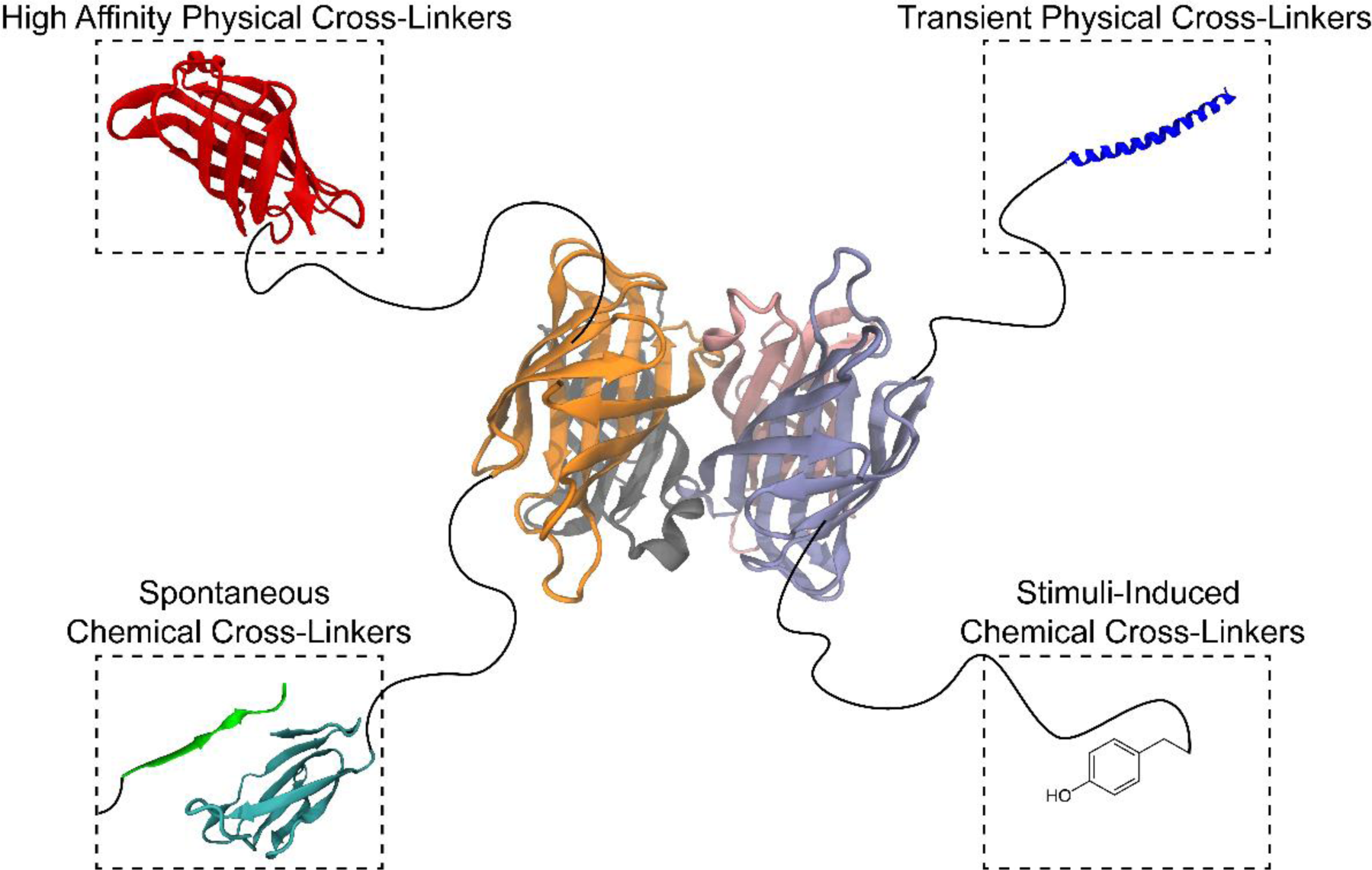
Schematic of the four-arm streptavidin precursors and second-level self-assembly building blocks for developing bottom-up design strategies that fabricate specific and stable protein-network materials. Streptavidin homotetramer (PDB: 1SWE) with customizable secondary crosslinking methods to control the mechanics of hydrogel fabrication. Secondary cross-linkers are illustrated together, attached to the same streptavidin cross-linker by a flexible protein strand, but were tested separately in this study to identify optimal mechanisms for the assembly of protein-network materials. Secondary cross-linkers, include a SAv monomer (SAv; red), coiled-coil protein monomer (P; blue; PDB: 1MZ9), SpyTag-SpyCatcher system (Spy; cyan and green; PDB: 4MLI), and tyrosine residue (Y) for photo-crosslinking. These results in artificial proteins of SAv-flexible protein-SAv, P-flexible protein-SAv, Spy-flexible protein-SAv and Y-flexible protein-SAv.

One distinct advantage of using SAv as a cross-linker is its ability to orthogonally control material properties. Beyond the tetra-SAv, we can go further by genetically fusing proteins of interest on both ends of SAv monomers, where the biosynthesized proteins will self-assemble into an eight-arm precursor (*f* = 8), octa-SAv. In addition, this molecular self-assembly approach still did not occupy the biotin binding site located in each SAv monomer, providing a mechanism for customizing octa-SAv precursors with the immobilization of biotinylated (macro)molecules, increasing up to 12 arm projections from an individual SAv cross-linker (8 ≤ *f* ≤ 12). Using recombinant protein technology to incorporate mechanical proteins as SAv arms that can form protein-network materials and attach biochemical (macro)molecules via biotinylation, will achieve the orthogonal control of mechanical and biochemical properties of the materials.

While the specificity and stability, as well as ligand-receptor property, of SAv open these possibilities to be used as multi-arm molecular nanohubs, the tetramer itself has not been used as a crosslinking mechanism to fabricate protein-network materials. Thus, whether the SAv stability at the molecular level can be transferrable to the bulk material level is unknown.

Here, we investigated the feasibility of using tetra-SAv as a protein-based alternative to tetra-PEG in the formation of protein-network hydrogels. To construct hydrogels, tetra-SAv-based nanomolecular building blocks were designed by the genetic fusion of a SAv monomer and a panel of biomolecular cross-linkers on either ends of well-characterized non-structured proteins (Figure 1). Four classes of biomolecular cross-linkers, such as SAv, coiled-coil, SpyTag-SpyCatcher, and dityrosine, were chosen to study the hierarchical network formation of four-arm SAv precursors. Unlike SAv, each type has been already used to prepare protein-based hydrogels^8, 15, 27^ and can be categorized as stable physical (i.e., non-covalent), transient (or weak) physical, specific and spontaneous chemical (e.g., covalent), and stimuli-induced chemical cross-linkers, respectively (Figure 1). The variability of the biomolecular cross-linkers enabled an accurate characterization of SAv cross-linkers, where non-covalently associated SAv cross-linkers in protein-network materials are stable and behave like chemical bonds with specificity. Simultaneously, the systematic analysis of nanomolecular building blocks from the bottom-up design of protein-network materials informs that the enhanced distribution of the building blocks within the network can be promoted by selecting the proper class of the secondary cross-linker. This, in turn, is crucial to reduce inhomogeneous crosslinking density and improve bulk mechanical properties of the protein-network material.^37, 38^ Collectively, the presented protein-network design that allows precise characterization of SAv cross-linkers provides protein assembly strategies for incorporating functionality, specificity, binding strength, and kinetics of cross-linkers to optimize material performances. Furthermore, this design approach is expected to impact the development of customizable material systems for diverse biotechnological applications, ranging from a synthetic scaffold that precisely mimics the extracellular microenvironment for tissue engineering^39, 40^ to electronic implantables.^41^

## Results and Discussion

### Artificial Protein with Tetra-SAv Cross-linkers

To analyze the properties of protein-network materials composed of tetra-SAv cross-linkers, we designed an artificial protein containing a SAv monomer on each end connected by an intrinsically disordered, flexible, polyelectrolyte-like protein containing 24 repeats of AGAGAGPEG amino acid sequence, denoted as C_24_,^27, 38^ which resulted in SAv-C_24_-SAv proteins (**Table S1**). We expected that SAv monomers in SAv-C_24_-SAv proteins would self-recognize and self-associate to form SAv homotetramers in aqueous buffer and develop hydrogels, similar to other well-known, non-covalently associating protein hydrogels.^14, 27, 38^ However, failed attempts to fabricate SAv-C_24_-SAv hydrogels between 10 – 20 w/v% led to an investigation of the binding mechanics. Here, we compared results of sodium dodecyl sulfate-polyacrylamide gel electrophoresis (SDS-PAGE) between SAv-C_24_-SAv samples that were denatured into individual, unstructured proteins by boiling at 100°C and samples that were not boiled (**Figure 2a**) as it is known that unboiled SAv maintain its tetramer structures and are observed in SDS-PAGE.^42^ Based on the two-fold difference in molecular weight between the boiled and unboiled conditions as well as the lack of bands at higher molecular weights in unboiled samples on the SDS-PAGE, we deduced that only two SAv-C_24_-SAv proteins are interacting to form tetramers without higher order binding that would indicate protein-network formation. The close proximity of two SAv cross-linkers connected by a flexible C_24_ strand allowed SAv on the N- and C-terminus to bind within each individual protein, forming a self-loop with the high specificity and affinity^43, 44^ (Figure 2b). Subsequently, two self-looped SAv-C_24_-SAv proteins interact with each other to form a tetrameric SAv complex. This binding sequence explains the lack of gelation and highlights the difficulty of forming hydrogels with cross-linkers that have high affinity and specificity connected by a flexible strand in an A-B-A triblock configuration.

**Figure 2.**
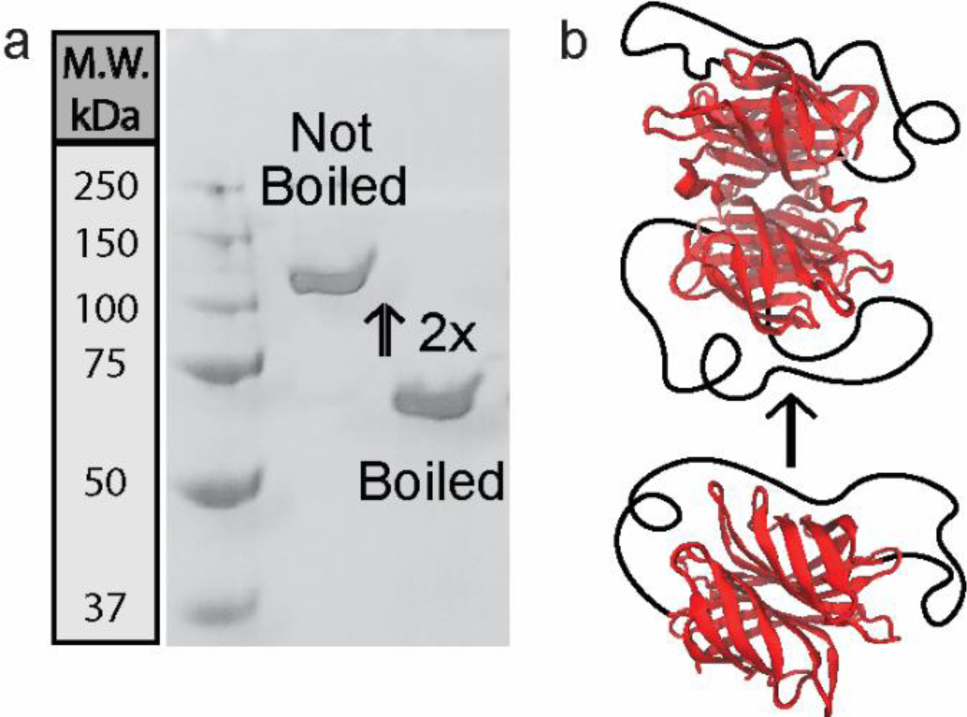
Analysis of SAv-C_24_-SAv homotetramerization. (a) SDS-PAGE analysis of SAv-C_24_-SAv self-association. (b) Schematic demonstrating the self-looping of an individual SAv-C_24_-SAv protein, followed by the molecular self-assembly of two self-looped SAv-C_24_-SAv proteins into a tetramer.

To prevent self-looping, we switched from the A-B-A polymer design, where streptavidin monomers are on both termini of the unstructured protein, to an A-B-C triblock configuration, where we limited our artificial protein designs to a single SAv cross-linker at the C-terminus and other physical or chemical crosslinking mechanisms at the N-terminus (Figure 1). With this design, SAv monomers form four-arm SAv precursors similar to the well-investigated tetra-PEG,^19, 21, 45^ can be engineered to initiate gelation, and exhibit mechanical stability with the help of the secondary cross-linkers. For the secondary crosslinking method, we initially considered typical protein chemical crosslinking technique that forms covalent bonds between chemical groups, such as amine and carboxyl groups, on either end of protein strands. Although strong and stable, these chemical bonds generate random crosslinking junctions because of the prevalence of chemical groups in the side chains of protein structures incorporated into network strands. The lack of the crosslinking specificity loses the control over key parameters that affect the mechanical properties of protein-network materials, such as chain length and network topology.^37^ To improve the specificity of this conventional chemical crosslinking strategy, molecular recognition (i.e., specificity) has developed prior to forming chemical bonds.^15, 25, 36, 46, 47^

### SAv-based Protein-Network Hydrogels with Secondary, Specific, Spontaneous, Chemical Cross-linkers

To apply the specific chemical crosslinking strategy, we implemented SpyTag-SpyCatcher chemistry, well-known for its ability to form spontaneous covalent isopeptide linkages between Asp117 of the SpyTag peptide and Lys31 of the SpyCatcher protein after the molecular recognition between those two molecules.^46^ We genetically incorporated either SpyTag or SpyCatcher to the N-terminus of C_24_-SAv and biosynthesized designed proteins (**Figure 3a-b**; Table S1). When mixing the solutions that contains either SpyTag-C_24_-SAv or SpyCatcher-C_24_-SAv, we observed a partially gelated slurry with dense gel formation where the two samples first interacted. However, a significant portion of the sample remains in liquid states due to the limited control of adequate mixing.^48^ This suggests that spontaneous crosslinking is susceptible to fabricating inhomogeneous protein-network materials. Protein– or polymer– network inhomogeneity is a debilitating issue in the field, diminishing mechanical properties due to inefficient gelation kinetics and requiring fabrication techniques that improve the homogeneity of the mixture during the network formation.^37, 48^

**Figure 3.**
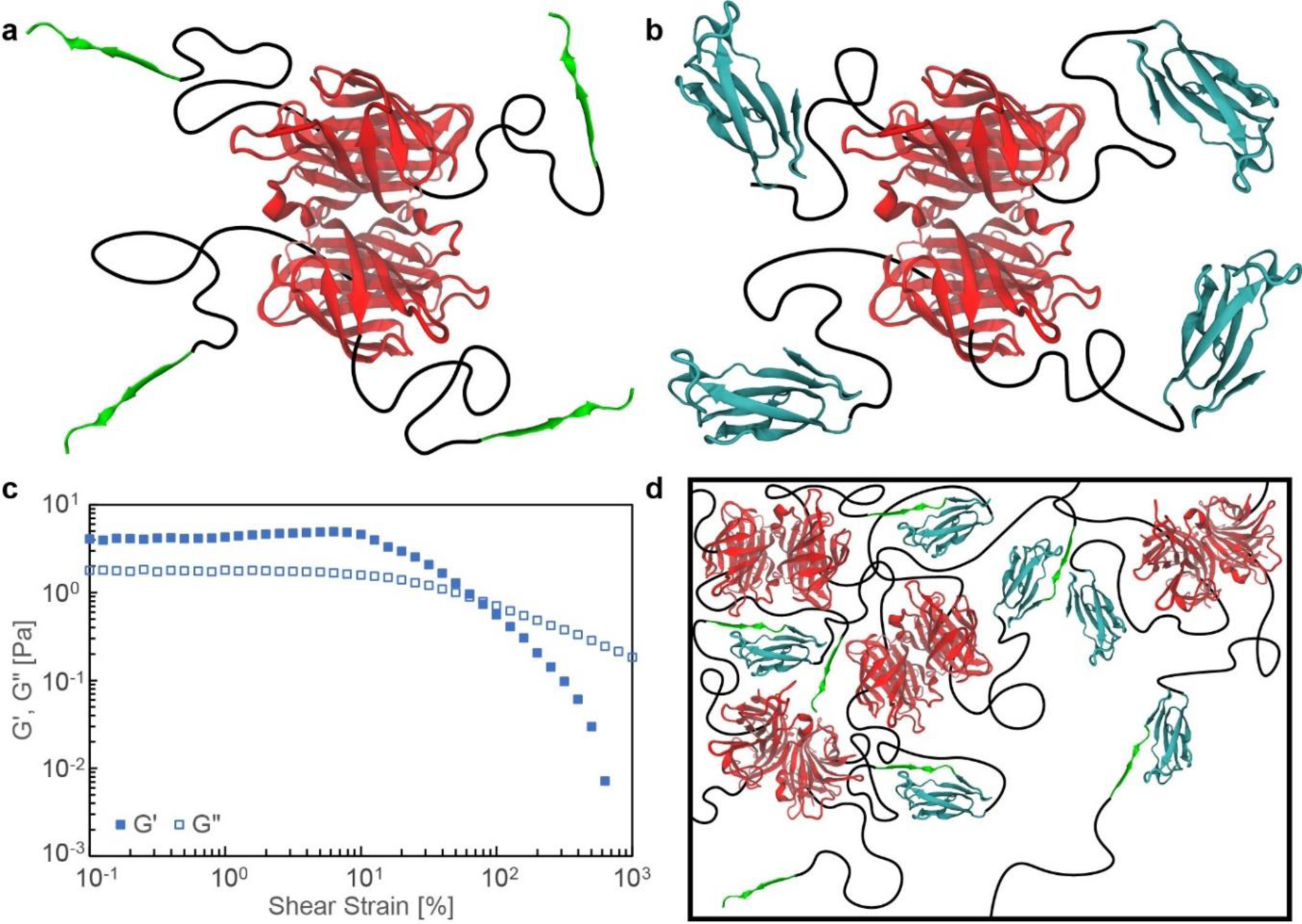
Spy-C_24_-SAv protein hydrogels and their rheological characterization. (a-b) Schemetics of a SpyTag-C_24_-SAv four-arm precursor and a SpyCatcher-C_24_-SAv four-arm precursor. (c) Representative strain sweep of a Spy-C_24_-SAv hydrogel at a constant 10 rad/s angular frequency. Spy denotes that the lingkage of SpyTag and SpyCatcher. (d) Illustration demonstrating the inhomogeneous protein-network formation of Spy-C_24_-SAv.

These qualitative observations were reflected in the rheological characterization (Figure 3c). Small amplitude oscillatory shear rheology captured the fast initial gelation from the Spy-C_24_-SAv network (**Figure S1**), where Spy represents the linkage between SpyTag and SpyCatcher. However, the material is relatively weak at less than 100 Pa of shear elastic modulus (G′; 0.02 ± 0.01 kPa; *N*=4). This measurement is within the 500 Pa range from prior studies that developed hydrogels using other SpyTag-SpyCatcher design strategies.^15, 47^ However, this G′ is weak relative to other protein-network materials with similar molecular weight and is significantly lower than the predicted G′ based on the polymer-network models^20, 38^ (**Table S2**). The experimental G′ is approximately 1% of the theoretical G′ from the affine and phantom polymer network models. As observed during the gelation, the specific and spontaneous covalent bonds seem to initiate protein-network formation prior to adequate diffusion of protein building blocks, resulting in an inhomogeneous crosslinking density within the protein network (Figure 3d) that limits the ability to characterize the mechanical properties of SAv cross-linkers.

### SAv-based Protein-Network Hydrogels with Secondary, Transient Cross-linkers

In addition to the specificity of biomolecular cross-linkers, their binding kinetics, dependent upon their binding affinity of non-covalent interactions,^49^ provide the ability to restructure the protein network and improve homogeneously distributed protein building blocks.^50, 51^ Biosynthetic coiled-coil protein structures, inspired by a variety of innate proteins such as cartilage oligomeric matrix protein and troponin, have been utilized as cross-linkers for the development of protein-network hydrogels with customizable network properties.^27, 28, 38, 50, 51^ Coiled-coil cross-linkers, where multiple ɑ-helical structures physically inter-wound each other, undergo transient binding kinetics where cross-linkers can bind and unbind periodically. These temporary bonds promote protein-network homogeneity by allowing protein building blocks to diffuse throughout the hydrogel over time.^49–51^ This ability to reorganize protein networks presented a viable option for improving network homogeneity in our tetra SAv-based protein-network design, engineering a monomer of pentameric coiled-coil cross-linker (P) at the N-terminus to form P-C_24_-SAv (**Figure 4a**; Table S1). SAv monomers in each protein led to form SAv tetramers with four protein arms (Figure 4a) due to its high affinity^23, 35^ as we confirmed on the SDS-PAGE^42^ (Figure 4d). Then, transient P cross-linkers are expected to construct protein-network hydrogels (Figure 4b). Upon adding physiological buffer, the artificial protein with 10 w/v% formed a hydrogel with good qualitative uniformity (Figure 4c), compared to the partial gelation of Spy-C_24_-SAv spontaneous crosslinking system, and showed no signs of protein aggregation as the hydrogel is optically transparent. During the SDS-PAGE analysis (Figure 2 and Figure 4d), we noticed that the protein samples ran slowly, which is caused by C_24_ as it was already reported by our previous work.^38^ We further confirmed its molecular weight using liquid chromatography-mass spectrometry (**Figure S2**). Characterizing the mechanical properties using rheology, we measured G′ of P-C_24_-SAv gels to be 1.6 ± 0.1 kPa (*N*=3; Figure 4e), a significant improvement compared to the Spy-C_24_-SAv hydrogels (0.02 ± 0.01 kPa; *N*=4; p < 0.01; see Methods Section). Also, the effective crosslinking density is improved from ∼1% to 26% and 47% when comparing the experimental G′ to the affine and phantom polymer network models (Table S2).

**Figure 4.**
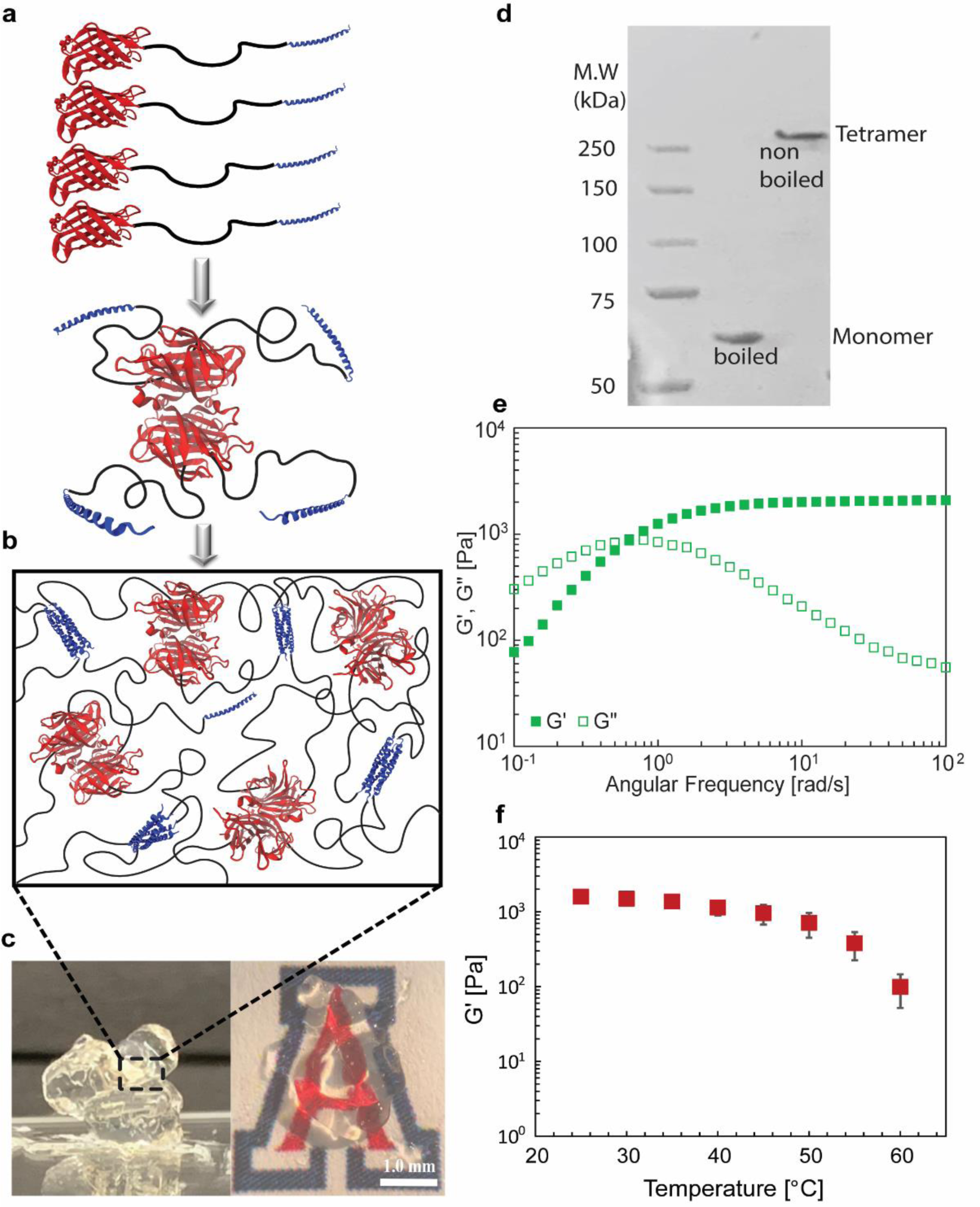
Characterization of SAv-based hydrogels with secondary, transient cross-linkers. (a) Schematic of a P-C_24_-SAv four-arm precursor from the molecular self-assembly of P-C_24_-SAv monomers. (b) Illustration demonstrating the protein-network formation of P-C_24_-SAv after the second molecular self-assembly of P coiled-coil cross-linkers. (c) Pictures of self-standing hydrogel and hydrogel under the light microscope. (d) SDS-PAGE analysis of boiled and non-boiled P-C_24_-SAv proteins. See the molecular weight of P-C_24_-SAv monomers, measured by liquid chromatography-mass spectroscopy (LC-MS) in Figure S2. (e) Representative frequency sweep of a P-C_24_-SAv hydrogel at a constant 1% strain. (f) Temperature sweep of P-C_24_-SAv hydrogels from 25°C to 60°C at a constant 10 rad/s frequency and 1% strain (*N*=3; mean ± standard error).

Even though the increase in G′ (Figure 4b) can correlate to improvements in the overall mechanical properties, P-C_24_-SAv hydrogel still lacked enough stability to characterize the properties of SAv cross-linkers. SAv tetramers are known to exhibit robust thermal stability that prevent their denaturation at temperatures up to approximately 85°C.^35^ During the rheological characterization of P-C_24_-SAv hydrogel, we increased the temperature to investigate whether this thermal stability of SAv tetramers translate into the bulk mechanical properties. G′ decreased by 93.9 ± 5.4 % (*N*=3) at 60°C compared to 25°C (Figure 4f). This could be due to the transient kinetics of P cross-linkers where higher temperatures promote the dissociation of coiled-coil structures.^50, 51^ However, it also cannot be excluded that the G′ drop can be due to the dissociation of non-covalently associated SAv homotetramers in the bulk hydrogel. Overall, the specific and transient physical associations of coiled-coil cross-linkers are beneficial because the transient kinetics can increase the building block homogeneity within protein networks (Figure 4d) compared to the specific and spontaneous covalent bonds of SpyTag-SpyCatcher cross-linkers (Figure 3). However, to adequately characterize the properties of tetra-SAv in protein-network hydrogels, the secondary crosslinking mechanism requires stability that exceeds the mechanical strength of SAv tetramers, in addition to the specificity and controlled binding kinetics.

### Mechanical and Thermal Stabilities of Tetra-SAv Cross-linkers in Protein-Network Hydrogels with Secondary, Stimuli-induced, Chemical Cross-linkers

Dityrosine photo-initiated chemical crosslinking can be the proper secondary crosslinking method because proteins with tyrosine residues can diffuse in solution prior to the formation of dityrosine covalent bonding between the tyrosine residues by blue-light irradiation. To avoid unintended dityrosine chemical crosslinking, the flexible C_24_ chain, which contains uncontrolled tyrosine residues, was replaced by elastin-like polypeptide (ELP) with a similar number of amino acid residues to C_24_ (Table S1). ELP has been extensively studied for its biocompatible and stimuli responsive properties for biomaterial applications.^52, 53^ It is also well-known for simple and effective purification of target proteins when genetically fused to ELP, using its temperature-responsive phase transition in solution.^54^ In our previous study, we used ELP with controlled numbers of tyrosine residues to modulate mechanical properties in photo-crosslinked protein-network materials.^55^ We adapted the ELP sequence and further engineered to prepare A_2_YA_2_-A_48_-SAv artificial proteins (**Figure 5a**), where A and Y represent the five amino acid sequence, VPGXG with either alanine (A) and tyrosine (Y) in the X position, respectively. Therefore, offering only one site of photo-crosslinking at the end of each ELP arm, which is at the opposite terminus to the SAv physical cross-linkers (Figure 5a; Table S1).

**Figure 5.**
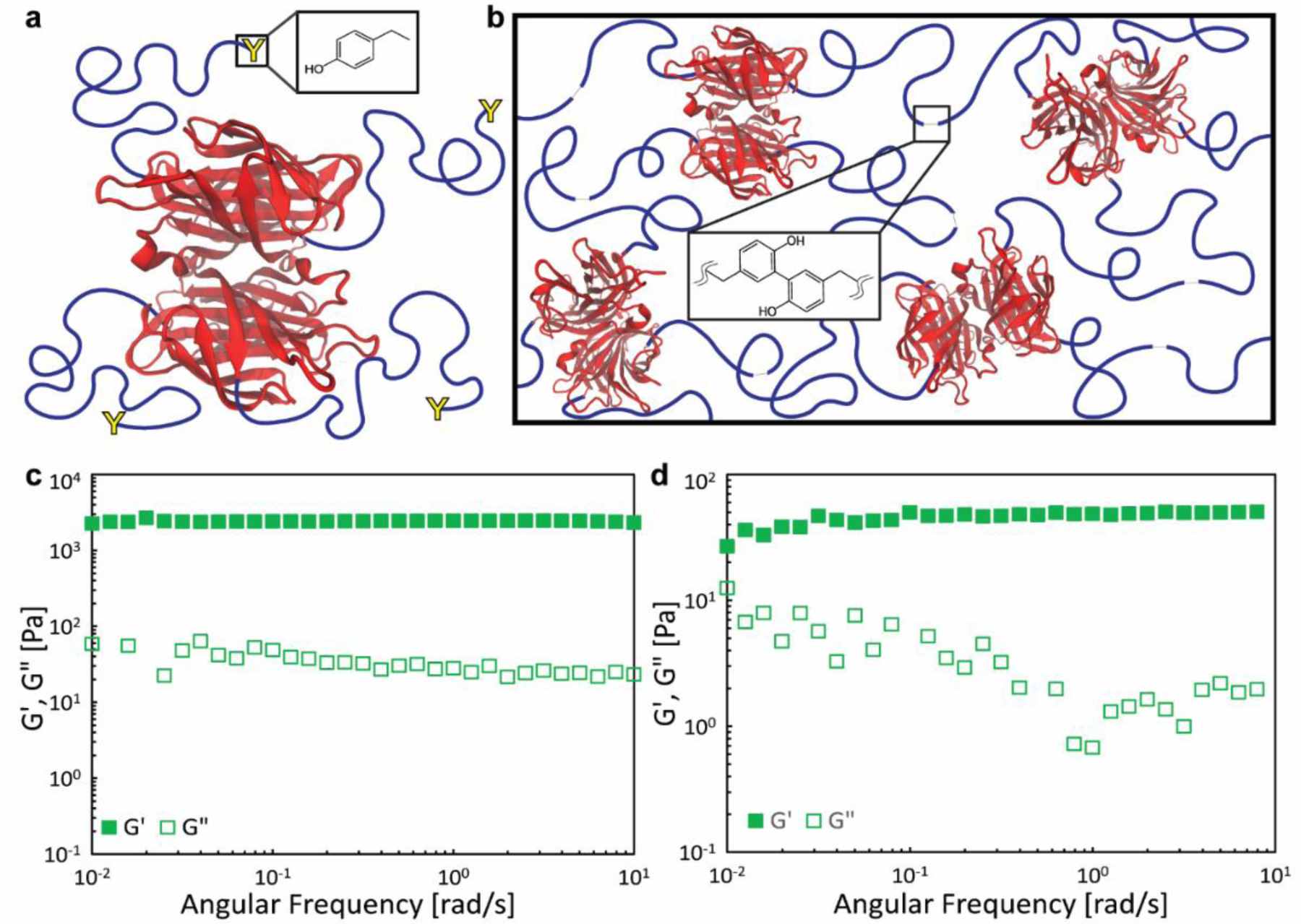
Mechanical stability of SAv-based hydrogels with secondary, photo-induced chemical cross-linkers using rheological characterization. (a) Tyr-ELP-SAv four-arm precursor. (b) Illustration demonstrating the protein-network of Tyr-ELP-SAv due to the formation of dityrosine covalent bonds and SAv tetramer self-assembly. (c-d) Representative frequency sweep of a Tyr-ELP-SAv hydrogel at a constant 1% strain. The proteins were photo-crosslinked with 250 µM ruthenium and 80 mM ammonium persulfate (c) and with 31 µM ruthenium and 15 mM ammonium persulfate (d).

The A_2_YA_2_-A_48_-SAv protein design successfully combined dityrosine chemical crosslinking and SAv physical crosslinking. The experimental G′ of the hydrogel, composed of four-arm SAv precursors with secondary dityrosine photo-crosslinking (Figure 5a-b), is 2.8 ± 0.4 kPa (*N*=3), which had an approximately 700-fold increase compared to the G′ of the Spy-C_24_-SAv hydrogel (p < 0.01; Figure 3c). In addition, the experimental G′ is approximately 70% of the theoretical G′ from the affine network model (Table S2). The experimental G′ was expected to be lower than the theoretical G′ since it is well-known that there are topological defects, such as self-loops and dangling chains, in polymer/protein networks with flexible strands.^20, 37, 38^ However, we see a significant improvement in hydrogel fabrication and G′ when replacing the SpyTag-SpyCatcher mechanism (*f* = 2) to a more controllable dityrosine photo-crosslinking (*f* = 2). This confirms that controlling the secondary crosslinking allows enough time for these cross-linkers to diffuse throughout the protein network before permanent chemical bond formation, which is crucial in improving mechanical properties or effective junctions in the network.

Interestingly, the G′ of A_2_YA_2_-A_48_-SAv hydrogel did not show the same frequency dependency as P-C_24_-SAv (Figure 4e), where G′ is dominating over G″ even at low frequencies despite the physical interactions of SAv tetramers (Figure 5c). This indicates that the reason why the P-C_24_-SAv acts like a physical gel, where the G′ of P-C_24_-SAv is frequency dependent, is because of the P cross-linker (Figure 4b). In contrast, the dityrosine photo-crosslinking mechanism causes the A_2_YA_2_-A_48_-SAv hydrogel to behave like a chemically crosslinked hydrogel although SAv tetramers are physically associated. Therefore, we confirm that SAv is a mechanically stable cross-linker with high specificity.

A caveat of dityrosine chemical crosslinking in protein-network materials is the limited control of the specificity in crosslinking when incorporating natural proteins where their structures require Tyr residues. Although we controlled the location and number of Tyr residues in the ELP sequence (Figure 5a), the SAv sequence contains Tyr residues that cannot be removed without possibly compromising its structure. These Tyr residues could generate unintentional SAv-A2YA2 or SAv-SAv dityrosine chemical crosslinking, potentially explaining the lack of frequency dependency in a non-covalently associated physical hydrogel (Figure 5c) and the experimental G′ approximately 140% of the theoretical G′ from the phantom network model^20^ (Table S2). However, there is a possibility that this unintentional dityrosine chemical crosslinking does not affect the crosslinking mechanism of SAv. The dimer-dimer interface of SAv homotetramers is the location where the SAv cross-linker will break by the external stress since that interface is weaker than the monomer-monomer interface.^23, 35^ In the case of SAv-A2YA2 crosslinking, the single tyrosine at the end of the ELP cannot covalently bind at the dimer-dimer interface due to the absence of tyrosine residues in that region (**Figure S3**). Therefore, this should not affect the physical interaction at the weak dimer-dimer interface which supports the fact that the lack of frequency dependency is not caused by any SAv-A2YA2 crosslinking.

The possibility of SAv-SAv crosslinking for the lack of frequency dependency (Figure 5c) was determined using SDS-PAGE by evaluating whether tyrosine residues in SAv can introduce unspecific dityrosine crosslinking. We found that photo-crosslinked SAv tetramers at 10 w/v% can form unspecific dityrosine bonds with the photo-crosslinking reagents of 250 µM ruthenium and 80 mM ammonium persulfate (**Figure S4**). However, the Tyr residues are not located at the dimer-dimer interface of SAv tetramers based on molecular visualization (Figure S3). This suggests that the non-covalently associated dimer-dimer interface of SAv cross-linkers are still intact and can be dissected during rheology characterization. Since we did not see the physical behavior in hydrogel rheology, this indicates that the physically associated SAv tetramer behaves like chemical cross-linker with the high specificity.

To confirm whether any SAv-SAv crosslinking affects the interfaces of SAv tetramers, we prepared the hydrogel with the reduced reagent concentration to 31µM ruthenium and 15mM ammonium persulfate which minimized unspecific crosslinking between SAv tetramers (Figure S4). While the reduced concentration impacted the G′ as expected based on our previous report on ELP gels,^55^ crossover frequency was not still observed (Figure 5d). Therefore, we concluded that, due to the high mechanical stability, the specifically self-associated, tetra-SAv in protein-network hydrogel behaves like a chemical cross-linker despite the fact that tetra-SAv is formed from non-covalent interactions.

To correlate the thermal stability of SAv from bulk, biochemical measurements to tetra-SAv within protein-network materials, the temperature-dependence of SAv-based hydrogels were evaluated. In temperature sweep rheology, we identified that the G′ of the hydrogel significantly increased at > 60°C (**Figure 6a**). This was caused by the incorporated ELP that has the lower critical solution temperature (LCST) behavior at approximately 60°C (Figure 6b) due to the phase separation or coacervation property of ELP which occur above certain temperature depending on ELP sequences. As we expected, this affect was shown in the changes of G′ across temperatures, where the intersection of the trendlines correlates with the occurrence of the ELP coacervation (Figure 6a). In addition, this mechanical reinforcement was already observed by the other ELP-based material system.^53^ When we further increase the temperature above the LCST and performed the frequency sweep experiment at 85°C (Figure 6c), we observed G′ is greater than G″ throughout angular frequencies below 10 rad/s where the crossover frequency (G′ = G″) or fluid behavior (G″ > G′) are generally present when characterizing physically-associated hydrogels (e.g., Figure 4e). Note that there were increases in the noise of the G″ data, meaning that the measurements were susceptible due to the decrease in concentrations of photo-crosslinking reagents (Figure 5d and Figure S4) which lowers the associated effective crosslinking density. It is also possible that higher temperatures than the ELP LCST (Figure 6c and Figure S4) potentially impacted the G″ due to the coacervation property of ELP. Nonetheless, we still clearly noticed that increasing temperature for Tyr-ELP-SAv hydrogel with low ruthenium and ammonium persulfate (Figure 5d) did not show the crossover frequency (**Figure S6**). As a result, the absence of a crossover frequency even at high temperatures and lower number of photo-crosslinked dityrosine bonds indicates the SAv tetramer was not responsible for any shear-stimulated, dissociation of its crosslinking when complemented with a secondary, transient cross-linker (Figure 4e). Furthermore, the data indicate that the tetra-SAv in protein-network materials is intact at the extreme temperature, confirming that tetra-SAv precursors are not only mechanically stable, but also thermally stable.

**Figure 6.**
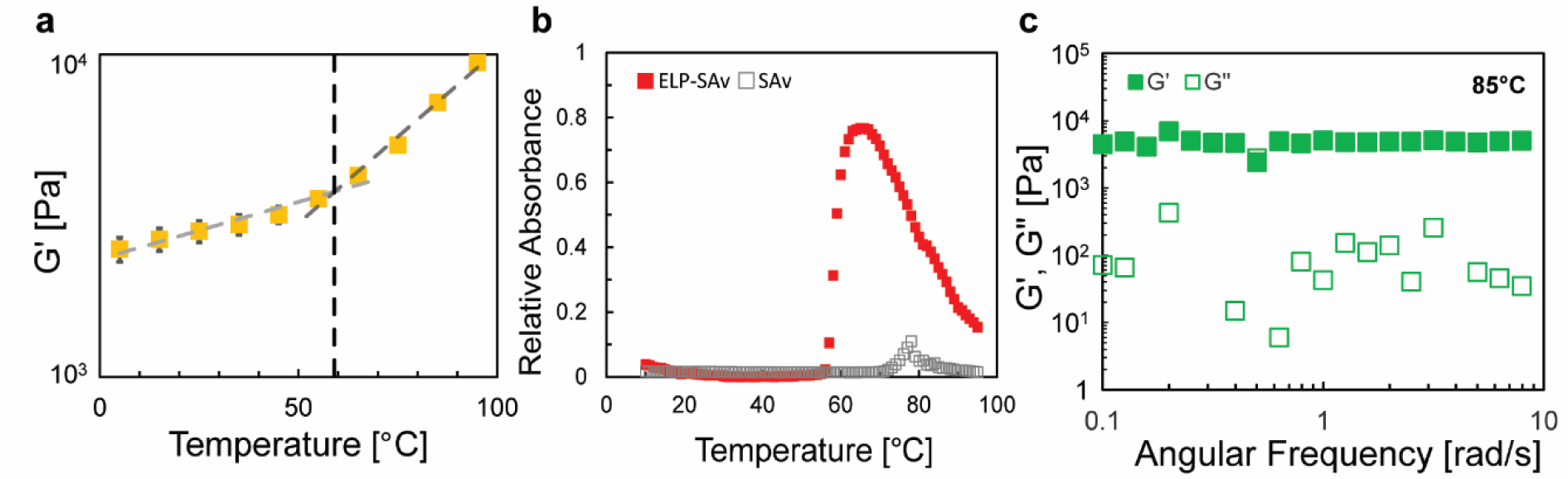
Thermal stability of tetra-SAv in protein-network hydrogels. (a) Temperature sweep rheology experiments on A_2_YA_2_-A_48_-SAv hydrogels with a constant 10 rad/s frequency and 1% strain. Trendlines were made from 5°C to 55°C (light gray dashed line) and from 65°C to 95°C (dark gray dashed line). Black dashed line represents the intersection. (b) UV-Vis measurement (at 350 nm) of A_2_YA_2_-A_48_-SAv proteins and SAv proteins as a control in the temperature ranges from 10°C to 95°C. UV-Vis data of SAv in wide range of wavelength at 25°C is in Figure S5. (c) Representative frequency sweep data of A_2_YA_2_-A_48_-SAv hydrogel at 85°C and a constant 1% strain.

While we successfully characterized the stability of SAv nanohubs in protein-network hydrogels, we identified that the secondary cross-linking mechanisms are critical on the effective crosslinking density. Stimuli-induced chemical crosslinking mechanism can potentially overcome a few topological defects in protein networks, such as self-loops (Figure 2) and inhomogeneous protein-network formation (Figure 3). Still, this system cannot be free from other topological defects, such as dangling chains, and higher-order loops,^37^ as experimental G′ is reduced compared to theoretical G′ (Figures 4-5; Table S2). By further modulating the flexibility of protein strands between biomolecular cross-linkers,^38^ this challenge can be addressed.

## Conclusion

Here, we investigated SAv tetramer as a stable and specific cross-linker for the development of protein-network materials or hydrogels. SAv was genetically engineered into artificial protein designs, resulting in individual proteins that self-recognize and self-associate with extreme affinity to form a four-arm tetramer hydrogel precursor. The systematic characterization of four-arm SAv-based hierarchical nanoassembly using a variety of second-level molecular self-assembly or specific chemical crosslinking methods discovered that stimuli-induced chemical cross-linkers, such as dityrosine photo-crosslinking, are optimal for developing protein-network materials with enhanced specificity, stability, and network homogeneity.

This customizable platform design can be engineered at the genetic level to incorporate target proteins with key mechanical, or biochemical properties in protein-network strands, or investigate alternative stimuli-induced chemical cross-linkers, such as GB1,^26^ to further improve secondary crosslinking specificity. Moreover, biotinylated molecules can be orthogonally coordinated with the SAv-based protein-network system by using the strong protein-ligand interaction of SAv-biotin for diverse biomedical applications, such as advanced cell signaling, drug delivery, biosensing, and imaging. Therefore, SAv cross-linkers as the molecular nanohub will be the essential component of functional protein-network materials.

## Materials and Methods

### Protein Expression and Purification

Target genes were cloned, expressed, and purified according to methods described in the referenced article.^38^ All gene sequences are described in Table S1. A_2_YA_2_-A_48_-SAv was purified using inverse transition cycling (ITC), exploiting the lower critical solution temperature (LCST) of elastin-like polypeptides.^54, 55^ After 4 cycles of ITC purification^55^, samples were dialyzed in deionized water to remove all salt impurities. All purified protein samples were flash frozen using liquid nitrogen, lyophilized, and stored at - 20°C.

### Physical Hydrogel Fabrication

Protein-based hydrogels with only physical cross-linkers, such as pentameric (P) coiled-coil and SAv, were prepared by hydrating lyophilized protein to 10% w/v in 20 mM tris buffer at pH 8.0.

### SpyTag-SpyCatcher Heterodimer Formation

Spy-C_24_-SAv hydrogels were prepared by hydrating lyophilized SpyTag-C_24_-SAv and SpyCatcher-C_24_-SAv proteins to 10% w/v, separately, in 20 mM tris buffer at pH 8.0, then mixing them at a 1:1 M ratio.

### Photo-crosslinking

Lyophilized A_2_YA_2_-A_48_-SAv proteins were dissolved in 20 mM tris buffer while incubating on a rocker at 4°C for approximately 1 hr. Ammonium persulfate and tris(2,2’-bipyridyl)ruthenium(II) chloride hexahydrate (Sigma-Aldrich, St. Louis, MO, USA) were dissolved in 20 mM tris and added to the protein solution for a final concentration of 15 or 80 µM and 31 or 250 mM, respectively, with 10% w/v A_2_YA_2_-A_48_-SAv. Hydrogels were prepared in a 4°C cold room by pipetting the mixture into a 14 mm diameter x 2 mm height mold and crosslinking 10 inches under a 24 W, 14 x 14 LED array emitting 460 nm blue light for 10 min. During photo-crosslinking, tris(2,2’-bipyridyl)ruthenium(II) ([Ru(II)bpy3]^2+^) oxidation of the tyrosyl phenyl groups form a spontaneous chemical bond, consuming a persulfate anion. Once completed, hydrogels were cut to an 8 mm diameter to match the rheometer geometry.

### Rheology

The viscoelastic mechanical properties were characterized using small amplitude oscillatory shear rheology on a Discovery Hybrid Rheometer 2 (TA Instruments, New Castle, DE, USA) with a sandblasted 20 mm cone-and-plate geometry at a 1° angle for physical secondary cross-linkers or a sandblasted 8 mm parallel plate geometry for chemical secondary cross-linkers and a sandblasted stage. Inertia, friction, and rotational mapping calibrations were performed prior to each experiment. A Peltier temperature-controlled stage maintained 25°C for all rheology testing, except during temperature sweeps. Hydrogels with physical secondary cross-linkers were transferred to the stage and the geometry was lowered to the trim gap height of 55 µm. Excess gel was removed from the edges of the geometry before lowering it to the testing gap height at 50 µm. Hydrogels with chemical secondary cross-linkers were pre-cut and loaded to an axial force of 0.05 N. To control evaporation, a mineral oil barrier was placed around the edges of the geometry and the geometry was encased in a solvent trap with a water seal. Hydrogels relaxed for 1 hour prior to experimentation. Strain sweeps were performed from 0.01 to 1000% shear strain at a constant 10 rad s^-1^ angular frequency. Frequency sweeps were performed from 0.01 to 100 rad s^-1^ at a constant 1% shear strain, within the linear region of the strain sweep. Depending on the type of hydrogel, temperature sweeps were performed between 5°C to 95°C, inclusively, at a constant 10 radˑs^-1^ angular frequency and 1% shear strain, within the linear region of the strain sweep.

### UV-Vis Temperature Sweep/Spectra

For ELP-SAv and only SAv proteins, absorbance at 350 nm were collected using a Cary 60 UV-vis spectrophotometer from 10 to 95°C using a Peltier temperature controller with an equilibration time of 5 minutes (Figure 6b). Absorbance spectra of SAv protein was collected between 270 and 800 nm at 25°C (Figure S5)

### Statistical Data Analysis

No data pre-processing or exclusion of outliers were applied when using Excel for statistical analysis. Rheology datasets pertaining to each hypothesis were analyzed using two-sided, independent samples t-tests to compare protein groups with a 0.01 threshold of significance. The sample size (*N*) for all experiments was *N* = 3, except for the Spy-C_24_-SAv system (*N* = 4). P-values for all t-test results were less than 0.01 and reported in the manuscript. All results written throughout the manuscript represent the mean ± standard deviation, unless otherwise noted.

### Theoretical Calculations

The theoretical shear elastic modulus (G′) of biopolymers was calculated using the affine and phantom network models (Table S2), according to methods described in detail in the referenced articles.^20, 38^

## ASSOCIATED CONTENT

### Supporting Information

The Supporting Information is available free of charge.

Illustrations of streptavidin tetramer with Tyr locations; Tyr crosslinking of SAv analyzed by SDS-PAGE; representative rheological characterization of protein hydrogels; full protein sequences; theoretical shear elastic modulus predictions based on network models. (PDF)

## Authors

**David S. Knoff** – *Department of Biomedical Engineering, University of Arizona, Tucson, AZ 85721, USA*

**Samuel Kim** – *Department of Biomedical Engineering, University of Arizona, Tucson, AZ 85721, USA*

**Kareen A. Fajardo Cortes** – *Department of Biomedical Engineering, University of Arizona, Tucson, AZ 85721, USA*

**Jocelyne Rivera** – *Department of Biomedical Engineering, University of Arizona, Tucson, AZ 85721, USA*

**Dallas Altamirano** – *Department of Biomedical Engineering, University of Arizona, Tucson, AZ 85721, USA*

## Notes

The authors declare no competing financial interests.

## ACKNOWLEDGMENT

We are grateful for our support from the National Heart, Lung, and Blood Institute of the National Institutes of Health (T32HL007955; D.S.K.), the ARCS® Foundation (D.S.K.), the Louise Foucar Marshall Foundation (D.S.K.), the Department of Education Federal TRIO Programs (Grant #P217A170284; K.A.F.C. and D.A.), NIH Maximizing Access to Research Careers grant (T34GM008718; J.R.) and the Tech Launch Arizona at the University of Arizona. The content is solely the responsibility of the authors and does not necessarily represent the official views of the National Institutes of Health.

