## Supplementary material for "Non-Covalently Associated Streptavidin Multi-Arm Nanohubs Exhibit Mechanical and Thermal Stability in Protein-Network Materials": SI

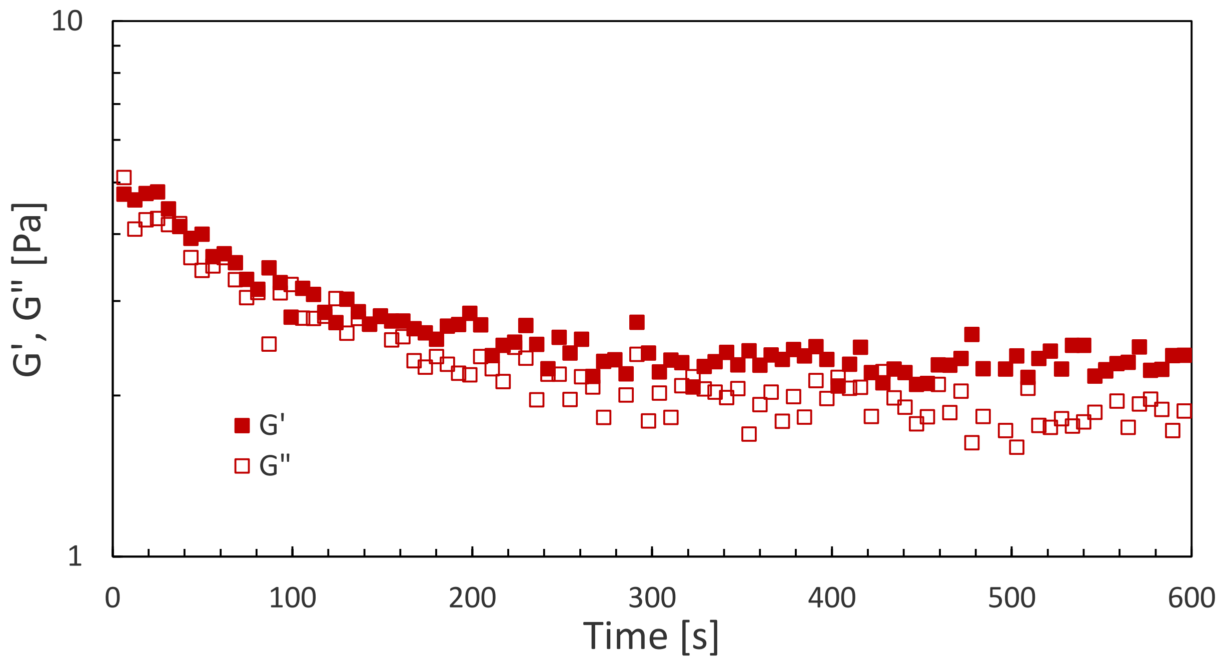


**Figure S1.** Time sweep rheology experiment capturing the gelation of Spy-$C_{24}$-SAv biopolymers.

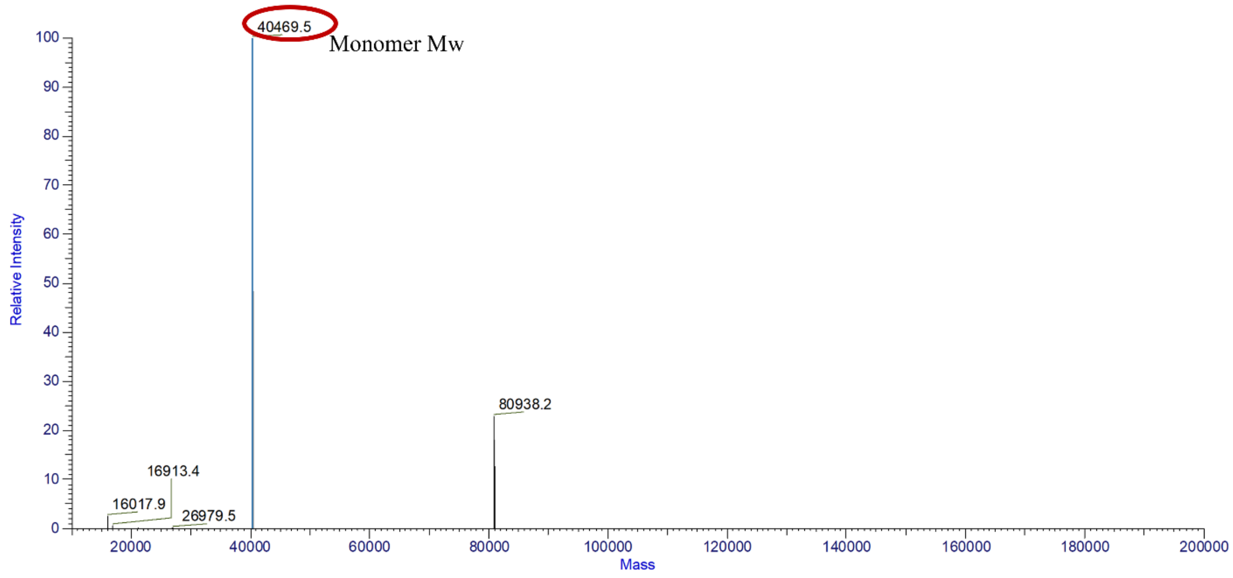


**Figure S2.** Liquid chromatography-mass spectrometry (LC-MS) of P-C24-SAv proteins (Figure 4). Dried samples were dissolved in 100% LC-MS-grade water. Ionized samples via electrospraying went through the vaporization process at 225℃ for a couple of seconds, which are common parameters for LC-MS analysis. Red circle indicates the detected M.W. of the protein monomer (~40.5 kDa). The vaporization process at the high temperature could denature or dissect P-C24-SAv tetramers (161.8 kDa) while some dimers (80.9 kDa) are observed, possibly due to the relatively strong monomer-monomer interaction, compared to the weak dimer-dimer interaction.[1]

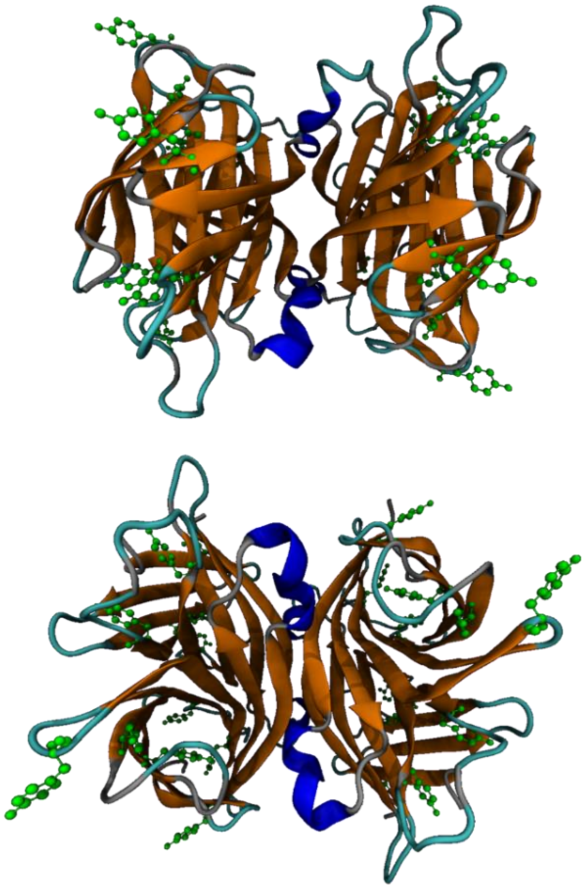

**Figure S3.** Two views of a streptavidin tetramer with tyrosine amino acids in green.

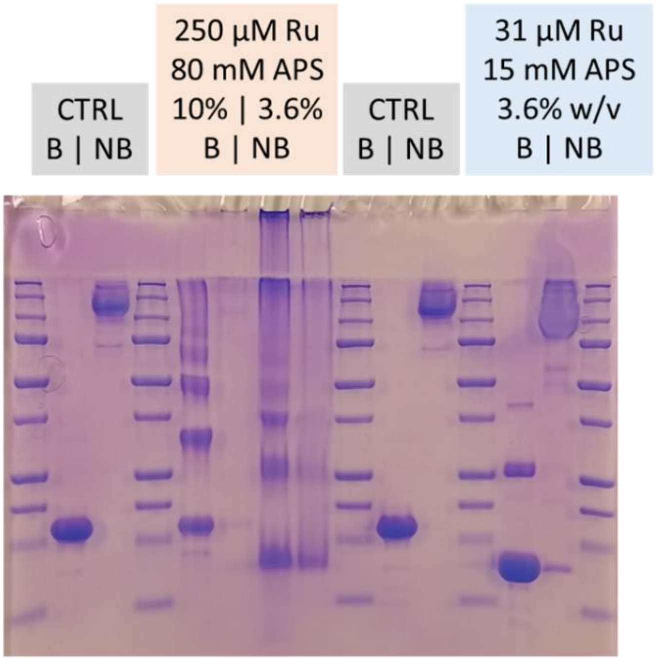

**Figure S4.** SDS-PAGE results show tetrameric streptavidin crosslinked using high and low concentrations of ruthenium and ammonium persulfate. To compare streptavidin tetramers and monomers, two samples were prepared for each combination of ruthenium and ammonium persulfate concentrations, where one sample (monomers) was boiled at 100℃ for 5 minutes and the other sample (tetramers) was left unboiled. For this experiment, streptavidin is obtained from Fisher Scientific (cat#: AAJ624288PL). Dityrosine photo-crosslinking method is described in the Experimental Section.

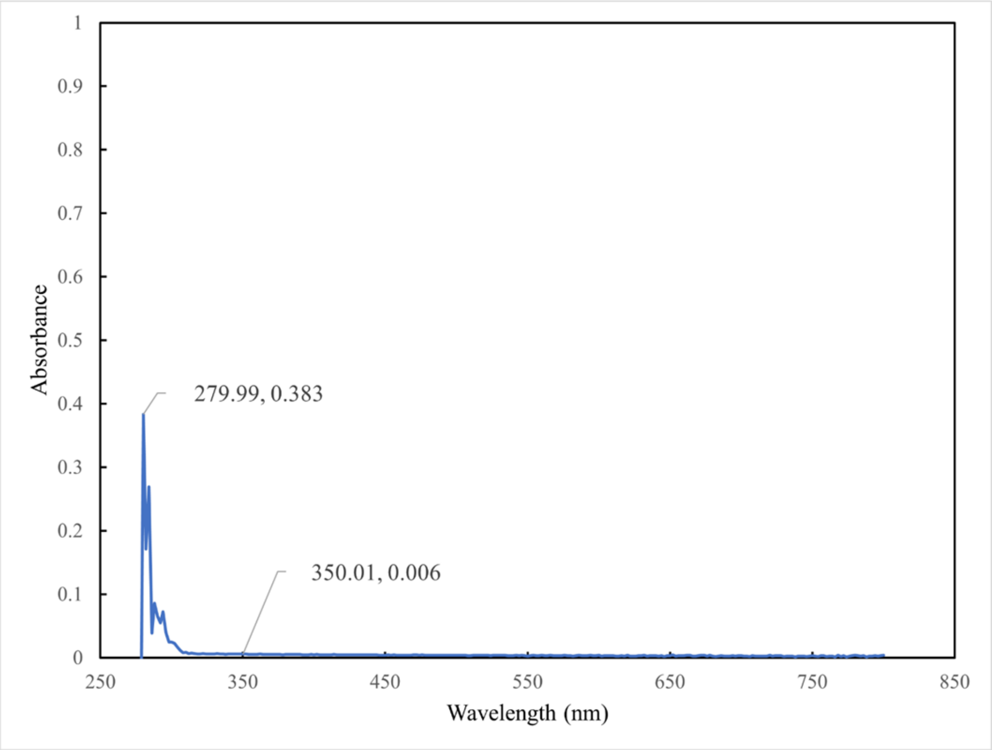


**Figure S5.** Absorbance spectra of SAv from 270 to 800 nm at 25℃. For this test, streptavidin was obtained from Fisher Scientific (cat#: AAJ624288PL).

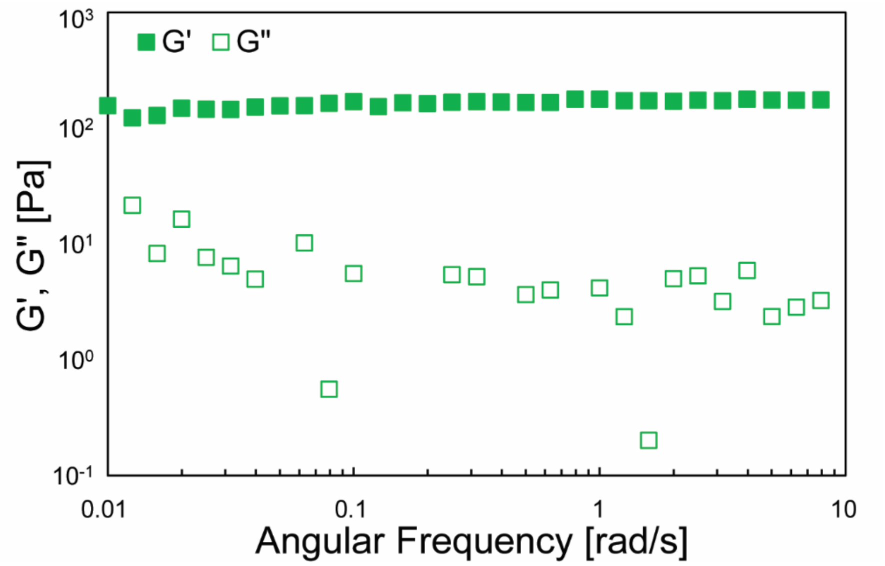


**Figure S6.** Rheological characterization data of a Tyr-ELP-SAv hydrogel photo-crosslinked with 31 µM ruthenium and 15 mM ammonium persulfate. Frequency sweep data was obtained at 75°C and a constant 1% strain. The same hydrogel tested at 4°C (Figure 5d) was used for this test.

**Table S1.** Full protein sequences for each protein studied. SAv cross-linkers (green highlight), P coiled-coil cross-linkers (underlined), $C_n$ (gray highlight), SpyTag (red highlight), SpyCatcher (purple highlight), ELP (turquoise highlight).

| Protein Name | Protein Sequence |
|---|---|
| SAv-$C_{24}$-SAv | *M*GSGSGSAEAGITGTWYNQLGSTFIVTAGADGALTGTYESAVGNAESRYVLTGRYDSAPATDGSGTALGWTVAWKNNYRNAHSATTWSGQYVGGAEARINTQWLLTSGTTEANAWKSTLVGHDTFTKVKPSAASASYRDPMGAGAGAGPEGAGAGAGPEGAGAGAGPEGAGAGAGPEGAGAGAGPEGAGAGAGPEGAGAGAGPEGAGAGAGPEGAGAGAGPEGARMPTSYRDPMGAGAGAGPEGAGAGAGPEGAGAGAGPEGAGAGAGPEGAGAGAGPEGAGAGAGPEGAGAGAGPEGAGAGAGPEGARMPTSYRDPMGAGAGAGPEGAGAGAGPEGAGAGAGPEGAGAGAGPEGARMPEFAEAGITGTWYNQLGSTFIVTAGADGALTGTYESAVGNAESRYVLTGRYDSAPATDGSGTALGWTVAWKNNYRNAHSATTWSGQYVGGAEARINTQWLLTSGTTEANAWKSTLVGHDTFTKVKPSAASKLAAALEHHHHHH |
| P-$C_{24}$-SAv | *M*GSGSGSAPQMLRELQETNAALQDVRELLRQQVKEITFLKNTVMESDASGASYRDPMGAGAGAGPEGAGAGAGPEGAGAGAGPEGAGAGAGPEGAGAGAGPEGAGAGAGPEGAGAGAGPEGAGAGAGPEGARMPTSYRDPMGAGAGAGPEGAGAGAGPEGAGAGAGPEGAGAGAGPEGAGAGAGPEGAGAGAGPEGAGAGAGPEGAGAGAGPEGARMPTSYRDPMGAGAGAGPEGAGAGAGPEGAGAGAGPEGAGAGAGPEGARMPEFAEAGITGTWYNQLGSTFIVTAGADGALTGTYESAVGNAESRYVLTGRYDSAPATDGSGTALGWTVAWKNNYRNAHSATTWSGQYVGGAEARINTQWLLTSGTTEANAWKSTLVGHDTFTKVKPSAASKLAAALEHHHHHH |

| | |
|---|---|
| SpyTag-$C_{24}$-SAv | *M*GSGSGSAHIVMVDAYKPTKASYRDPMGAGAGAGPEGAGAGAGPEGAGAG<br>AGPEGAGAGAGPEGAGAGAGPEGAGAGAGPEGAGAGAGPEGAGAGAGPEG<br>AGAGAGPEGAGAGAGPEGARMPTSYRDPMGAGAGAGPEGAGAGAGPEGAG<br>AGAGPEGAGAGAGPEGAGAGAGPEGAGAGAGPEGAGAGAGPEGAGAGAGP<br>EGAGAGAGPEGAGAGAGPEGARMPTSYRDPMGAGAGAGPEGAGAGAGPEG<br>AGAGAGPEGAGAGAGPEGARMPEFAEAGITGTWYNQLGSTFIVTAGADGA<br>LTGTYESAVGNAESRYVLTGRYDSAPATDGSGTALGWTVAWKNNYRNAHS<br>ATTWSGQYVGGAEARINTQWLLTSGTTEANAWKSTLVGHDTFTKVKPSAA<br>SKLAAALEHHHHHH |
| SpyCatcher-$C_{24}$-SAv | *M*GSGSGSAMVDTLSGLSSEQGQSGDMTIEEDSATHIKFSKRDEDGKELAG<br>ATMELRDSSGKTISTWISDGQVKDFYLYPGKYTFVETAAPDGYEVATAIT<br>FTVNEQGQVTVNGKATKGDAHIDASYRDPMGAGAGAGPEGAGAGAGPEGA<br>GAGAGPEGAGAGAGPEGAGAGAGPEGAGAGAGPEGAGAGAGPEGAGAGAG<br>PEGAGAGAGPEGAGAGAGPEGARMPTSYRDPMGAGAGAGPEGAGAGAGPE<br>GAGAGAGPEGAGAGAGPEGAGAGAGPEGAGAGAGPEGAGAGAGPEGAGAG<br>AGPEGAGAGAGPEGAGAGAGPEGARMPTSYRDPMGAGAGAGPEGAGAGAG<br>PEGAGAGAGPEGAGAGAGPEGARMPEFAEAGITGTWYNQLGSTFIVTAGA<br>DGALTGTYESAVGNAESRYVLTGRYDSAPATDGSGTALGWTVAWKNNYRN<br>AHSATTWSGQYVGGAEARINTQWLLTSGTTEANAWKSTLVGHDTFTKVKP<br>SAASKLAAALEHHHHHH |
| $A_2YA_2$-$A_{48}$-SAv | *M*HMVPGAGVPGAGVPGYGVPGAGVPGAGPWVPGAGVPGAGVPGAGVPGAG<br>VPGAGVPGAGVPGAGVPGAGVPGAGVPGAGVPGAGVPGAGVPGAGVPGAG<br>VPGAGVPGAGVPGAGVPGAGVPGAGVPGAGVPGAGVPGAGVPGAGVPGAG<br>VPGAGVPGAGVPGAGVPGAGVPGAGVPGAGVPGAGVPGAGVPGAGVPGAG<br>VPGAGVPGAGVPGAGVPGAGVPGAGVPGAGVPGAGVPGAGVPGAGVPGAG<br>VPGAGVPGAGVPGAGVPGAGASAEAGITGTWYNQLGSTFIVTAGADGALT<br>GTYESAVGNAESRYVLTGRYDSAPATDGSGTALGWTVAWKNNYRNAHSAT<br>TWSGQYVGGAEARINTQWLLTSGTTEANAWKSTLVGHDTFTKVKPSAASL<br>EHHHHHH |

**Table S2.** Theoretical shear elastic moduli (G′) predictions based on Affine and Phantom network models. The methods for the calculation can be found in the referenced articles.[2,3]

| Cross-linker functionality | Biomolecular cross-linker | Molecular weight of protein between cross-linkers | ν ($m^{-3}$) | *Affine G′ (kPa) | Phantom G′ (kPa) |
|---|---|---|---|---|---|
| f=4 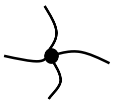 | Streptavidin tetramer | 46.89 kDa; SAv-$C_{24}$-SAv (Figure 2) | $1.30 \times 10^{24}$ | 5.30 | 2.60 |
| | | 82.58 kDa; SAv-$C_{24}$-SpyTag-SpyCatcher-$C_{24}$-SAv (Figure 3) | $7.30 \times 10^{23}$ | 3.00 | 1.50 |
| | | 68.02 kDa; SAv-ELP ($A_{48}$-$A_2YA_2$- – $A_2YA_2$-$A_{48}$-SAv (Figures 5-6) | $8.90 \times 10^{23}$ | 3.60 | 1.80 |
| f=4 and 5 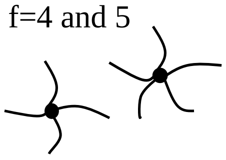 | Streptavidin tetramer & P coiled-coil pentamer | 40.47 kDa; P-$C_{24}$-SAv (Figure 4) | $1.50 \times 10^{24}$ | 6.10 | **3.40 |

* Affine G′ is independent from the cross-linker functionality.

** To calculate the P-$C_{24}$-SAv protein network's Phantom G′, we assumed that this theoretical G′ is between the two phantom G′ when f = 4 and f = 5, individually, since both functionalities are found in this protein network. To estimate the theoretical G′, we calculated the % ratio of SAv tetramer (f = 4) and P pentamer (f = 5) in terms of moles, which is 56% and 44%, respectively. Then, we summed up the product of each percentage to its respective phantom G′ (3.10 kPa when f = 4 and 3.70 kPa when f = 5). This led to an estimated theoretical G′ of approximately 3.40 kPa.
